# Nonenzymatic template-directed replication using 2′-3′ cyclic nucleotides under wet-dry cycles

**DOI:** 10.1101/2022.07.11.499554

**Authors:** Shikha Dagar, Susovan Sarkar, Sudha Rajamani

## Abstract

‘RNA World Hypothesis’ is centred around the idea of a period in the early history of life’s origin, wherein nonenzymatic oligomerization and replication of RNA resulted in functional ribozymes. Previous studies in this endeavour have demonstrated template-directed replication using chemically modified nucleotides and primers. Nonetheless, similar studies that used non-activated nucleotides led to the formation of RNA only with abasic sites. In this study, we report template-directed replication with prebiotically relevant cyclic nucleotides, under dehydration-rehydration (DH-RH) cycles occurring at high temperature (90°C) and alkaline conditions (pH 8). 2′–3′ cyclic nucleoside monophosphates (cNMP) resulted in primer extension, while 3′–5′ cNMP failed to do so. Intact extension of up to two nucleotide additions was observed with both canonical hydroxy-terminated (OH-primer) and activated amino-terminated (NH_2_-primer) primers. We demonstrate primer extension reactions using both purine and pyrimidine 2′–3′ cNMPs, with higher product yield observed during cAMP additions. Further, the presence of lipid was observed to significantly enhance the extended product in cCMP reactions. In all, our study provides a proof-of-concept for nonenzymatic replication of RNA, using intrinsically activated prebiotically relevant cyclic nucleotides as monomers.

## Introduction

Contemporary biology relies on dedicated biopolymers such as DNA/RNA to store and propagate genetic information, and proteins that enable the catalysis of reactions (1). As primitive cells are considered to be devoid of a refined and evolved molecular machinery, these functions might have been carried out by prebiotically relevant, simple molecular systems. Towards this, the well-explored ‘RNA World Hypothesis’ suggests RNA as having been the first biopolymer to have emerged due to its ability to serve both as a catalyst and a genetic material (2). In the absence of enzymes, the formation and replication of RNA would have had to be nonenzymatic; driven by both the monomeric species involved and the environmental conditions. Previous attempts to demonstrate the enzyme-free replication of RNA either involved chemically modified primer (3′-dideoxy-NH_2_ modified primer (NH_2_-primer)), nucleotides (imidazole, methylimidazole or aminoimidazole at 5′ end of the nucleotide), or both, and were performed at ambient temperature (24°C) (3–11). Although some studies have shown a plausible route for the formation of such activated nucleotides under prebiotic settings (12–16), their presence in significant amounts on the early Earth is questionable. Pertinently, their susceptibility towards hydrolysis at high temperature and in aqueous conditions, renders them as unsuitable substrates, putting into question their use as prebiotically compatible reactants.

In a relevant study, Bapat *et. al*. investigated phospholipid-assisted template-directed NH_2_-primer extension reactions using non-activated nucleotides i.e., nucleoside 5′-monophosphate (5′-NMP), under acidic hydrothermal conditions (17). The presence of amino group in the primer makes it a better nucleophile as compared to hydroxyl-terminated primer, facilitating its attack on the phosphorus group of the incoming nucleotide, resulting in the formation of a phosphoramidite bond. These reactions were performed under alternating dehydration-rehydration (DH-RH) cycles at high temperature (90°C) and acidic pH (pH 2). DH-RH cycles are a recurring geological phenomenon facilitated readily in terrestrial hydrothermal pools and have been demonstrated to facilitate condensation reactions and the formation of different biopolymers under prebiotic settings (18, 19). The dry phase concentrates the substrate molecules and the rehydrated phase facilitates mixing, thus, increasing the chances of effective collisions and redistribution of the growing oligomers. Lipids have been shown to form multilamellar films during the dry phase, which upon rehydration can encapsulate the surrounding solutes (e.g. informational oligomers), resulting in protocell-like entities (20– 22). However, similar to the oligomerization reactions, the systematic characterization of the primer extension product obtained under these reaction conditions (high temperature and pH 2), showed depurination of potentially the incoming nucleotide that resulted in loss of the information moiety (17, 23, 24). Importantly, similar nonenzymatic template-directed primer extension reactions with 5′-NMPs and hydroxyl-terminated primer (OH-primer), resulted in no prominent primer extension (17). This emphasizes the necessity of using prebiotically relevant alternative monomers, which do not require acidic pH for the formation of a phosphodiester bond, thereby alleviating the problem of the loss of information moiety.

Recently, we demonstrated the nonenzymatic oligomerization of prebiotically relevant cyclic nucleoside monophosphates (cNMPs) under DH-RH cycles that yielded intact RNA oligomers (25). Their prebiotically possible synthesis, stability to high temperatures, and being intrinsically active due to the presence of an intramolecular cyclic group, make cNMPs a potential substrate for nonenzymatic oligomerization and replication reactions (12, 13, 26–31). However, to our knowledge, no study has been reported wherein template-directed replication (primer extension) has been attempted using cNMPs. Therefore, we investigated the use of cNMPs in template-directed primer extension reactions of RNA. The two isomers of cNMP (2′-3′cNMP or 3′-5′cNMP) are known to possess different stability towards hydrolysis, and hence differ in their reactivity and oligomerization potential (25). Given this, the template-directed primer extension reactions were performed with both the structural isomers i.e., 2′-3′ cNMP and 3′-5′ cNMP using a purine, (cAMP) or a pyrimidine (cCMP) as the incoming monomer. The reactions were undertaken using both the OH-primer and NH_2_-primer (latter has been used extensively in many previous studies). As lipids have been shown to impart protection to RNA against hydrolysis under DH-RH conditions (17, 23, 32), the effect of the presence of lipids was also evaluated in these reactions. Previous studies that evaluated lipid-assisted template-directed replication have predominantly employed pure phospholipid (PL) membranes (17, 33). Although insightful, such membranes are impermeable to polar molecules and would require complex membrane proteins to facilitate such processes (34). The exchange of material with the surroundings is an essential feature of robust protocells, thus, making it implausible for just pure PL membranes to act as prebiotic compartments. In this regard, few other studies have investigated fatty acid/PL blended (hybrid) membranes as model protocellular compartments (35). Presence of a single chain amphiphile (SCA) like fatty acids in these blended membranes, enhanced the permeability of these compartments, while the use of a diacyl chain amphiphile like PL, enabled maintaining their stability against environmental fluctuations. Given this, we evaluated the effect of lipids on nonenzymatic primer extension reactions using three different model protocellular membranes i.e., only palmitoyl-2-oleoyl-sn-glycero-3-phosphocholine (POPC), and two binary systems of POPC with glycerol 1-monooleate (GMO) and POPC with oleic acid; both present in 1:1 molar ratio in the binary systems.

Our results showed the extension of the primer with two intact nucleotide additions, in the reactions involving 2′-3′ cNMPs at pH 8 and under DH-RH conditions. No significant primer extension was observed in the reactions involving 3′-5′ cNMPs. Importantly, the primer extension by two intact nucleotides was observed for both the OH-primer and NH_2_-primer, which would have resulted in phosphodiester and phosphoramidite linkage, respectively. The primer extension was observed with both 2′-3′ cAMP as well as 2′-3′ cCMP. Pertinently, in the 2′-3′ cCMP reactions, the presence of lipids was observed to significantly increase the yield of the extended product. Altogether, this study provides a comprehensive approach by which enzyme-free information propagation might have been facilitated on the early Earth in a putative RNA World that involves the use of prebiotically realistic, ‘intrinsically active’ monomers.

## Materials and methods

The monosodium salts of all four cyclic monophosphates viz. Adenosine 3′, 5′ cyclic monophosphate (3′, 5′ cAMP), Adenosine 2′, 3′ cyclic monophosphate (2′, 3′ cAMP), Cytidine 3′, 5′ cyclic monophosphate (3′, 5′ cCMP) and Cytidine 2′, 3′ cyclic monophosphate (2′, 3′ cCMP) were purchased from Sigma-Aldrich (Bangalore, India) and used without further purification. 1-palmitoyl-2-oleoyl-sn-glycero-3-phosphocholine (POPC) was purchased from Avanti Polar Lipids Inc (Alabaster, AL, USA); other co-surfactants i.e., glycerol 1-monooleate (GMO) and oleic acid (OA) were purchased from Nu-Chek-Prep (Elysian, MN, USA) and all were used without further purification. All other reagents used were of analytical grade and purchased from Sigma-Aldrich (Bangalore, India). The RNA primers used in this study are Amino-G (NH_2_-primer; acquired from Keck laboratory, Yale, USA), Hydroxyl-G (OH-primer) and 10-mer hydroxyl-G (10-mer OH-primer) (acquired from Thermo Fisher Scientific, USA) primers. Both Amino-G and Hydroxyl-G primers were fluorescently labelled with Cyanine 3 (Cy3) on their 5′-end for facilitating their detection on polyacrylamide gel electrophoresis (PAGE). The Amino-G primer terminates with a 3′-amino-2′, 3′-dideoxynucleotide (Metkinen, Finland) while the Hydroxyl-G primer terminates with a canonical ribonucleotide. The sequences of the primers and templates are as given below, with the template base indicated in bold:

Primer Amino-G (Amino/ NH_2_-primer): 5′ GG GAU UAA UAC GAC UCA CUG_NH2_-3′

Primer Hydroxyl-G (Hydroxyl/ OH-primer): 5′ GG GAU UAA UAC GAC UCA CUG_OH_-3′

10-mer primer hydroxyl-G (10-mer OH-primer): 5′ C GAC UCA CUG_OH_-3′

Template U: 5′ AGU GAU CU**U** CAG UGA GUC GUA UUA AUC CC_OH_-3′

Template G: 5′ AGU GAU CU**G** CAG UGA GUC GUA UUA AUC CC_OH_-3′

## Methods

### Reaction setup

In a typical reaction, 2.5 µM of the template (U in the case of cAMP and G in the case of cCMP) was annealed with 1.25 µM of the Cy3 labelled primer (either OH-primer or NH_2_-primer), by heating them at 95 °C for 5 minutes and then cooling to room temperature (RT). 5 mM of the corresponding cyclic nucleotide monophosphate (cNMP) was added to the annealed primer-template complex. 1 mM of either pure POPC or binary suspension of POPC:GMO::1:1 or POPC:OA::1:1 was added in case of the lipid-assisted reactions. The pH of the mixture was adjusted to ∼ 8, unless specified. The reaction mixtures were then subjected to repeated cycles of Dehydration-Rehydration (DH-RH) at 90°C, using 24 hours per cycle. After each dehydration cycle, the reaction mixture was rehydrated with 1 mM of the respective cNMP to compensate for potential loss of substrate. The samples were withdrawn after the rehydration step for analysis after regular intervals (i.e., starting of the reaction, after one day, two days and five days, respectively). These samples were then analyzed using denaturing PAGE.

### PAGE analysis of the reaction products

After varying time periods (i.e., starting of reaction, after one day, two days and five days), the samples were collected in 1X TBE buffer containing 8 M urea. The sample volumes withdrawn at different time points were adjusted to a standardized amount to compensate for the degradation of RNA primer that occurs over multiple cycles of DH-RH. A non-fluorescent competitor RNA (untagged primer), whose sequence was exactly the same as that of the tagged primer, was added in at least 10 times excess as compared to the fluorescently tagged primer, in all the samples that were to be run on the PAGE gel. This was done in order to successfully separate the fluorescent primer and the extended product from the template for unhindered and effective gel analysis. The reaction products were analyzed on 20% denaturing PAGE. The gels were then scanned with an Amersham Typhoon Biomolecular imager (GE Healthcare) at 550 PMT and 100-micron resolution setting, using the Cy3 (532 nm) excitation laser. The gel images were subsequently processed using ImageQuant v8.2 software for quantification of the relevant bands.

### Quantification of the reaction products

The yield (%) of the extended products of the primer was calculated by using the following formula:

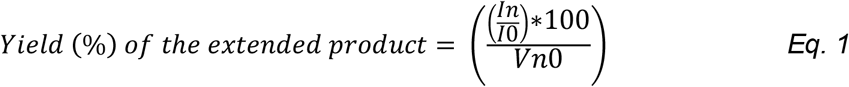

Where, I_n_ and I_0_ are the intensities of the band at the n^th^ cycle and the 0^th^ cycle, respectively, and V_n0_ is the volume fold change in n^th^ cycle with respect to 0^th^ cycle, calculated by:

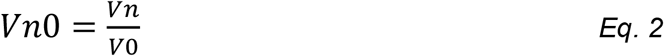

### LC-MS analysis of the extended products

For LC-MS analysis, primer extension reactions were performed using the 10-mer OH-primer. This was done in order to detect the undegraded extended primer under the conditions optimal for LC-MS analysis, as the longer the RNA, the higher is the possibility of its degradation upon ionization. A mixture of 1.25 µM of 10-mer primer and 5 mM of 2′, 3′ cNMP was subjected to DH-RH cycles at pH 8. After each dehydration cycle (24 hours per cycle), the reaction mixture was rehydrated with 1 mM of the respective cNMP. After two DH-RH cycles, all the reaction mixture timepoints from a specific reaction were pooled together (total volume 5 ml), in order to increase the total extended product to be present in detectable amount. This mixture was then lyophilized and resuspended in 52 µl, out of which 50 µl was loaded onto LC-MS. The reaction mixtures were separated on a Zorbax C8 column (dimensions: 4.6 × 150 mm, particle size: 3.6 µm) (Thermo Scientific), fitted with a guard column. The species were separated on LC using a gradient of solvent A of 95:5 (vol/vol) H_2_O/methanol (MeOH) + 0.1% ammonium hydroxide, and solvent B of 60:35:5 (vol/vol) isopropanol/MeOH/H_2_O + 0.1% ammonium hydroxide. The gradient involved an isocratic phase of solvent A for 10 minutes, followed by an increase to 100% of solvent B for six minutes and subsequent equilibration with solvent A for seven minutes. Mass spectra of the reaction samples was recorded on a Sciex X500R QTOF mass spectrometer (MS) fitted with an Exion-LC series UPLC (Sciex, CA, USA) using information-dependent acquisition (IDA) scanning method. The acquired data was analyzed using the Sciex OS software. All the mass acquisition was performed using Electron spray ionization (ESI) in the negative mode with the following parameters: turbo spray ion source, medium collision gas, curtain gas = 30 L/min, ion spray voltage = -4500 V (negative mode), at 500 °C. TOF-MS acquisition was done using a declustering potential of -80 V, and -10 V collision energy. As the mass acquisition was carried out in the negative mode, the observed masses correspond to the mass of the nH^-^ (where n is the number of deprotonation sites) adducts of the parent molecule. The presence of a specific species/molecule was confirmed by the presence of precursor mass within 10 ppm error.

## Results

### Primer extension using NH_2_-primer and cyclic nucleotides under DH-RH conditions

Given that cNMPs are prebiotically relevant nucleotides and can oligomerize nonenzymatically (25, 27, 28, 36, 37), we asked whether these can act as a substrate for enzyme-free RNA copying. Firstly, we investigated template-directed primer extension using an amino terminated primer (NH_2_-primer), with 2′, 3′ cAMP as the incoming nucleotide, under multiple DH-RH cycles at 90°C and varying pH (2, 8 and 10) (Fig. 1a and S3). As alluded to earlier, the rehydrated reaction mixture was withdrawn after various time periods and analyzed using denaturing PAGE (detailed in the Methods section). A reaction with known single nucleotide addition to the original 20-mer amino primer was used as the control (indicated by ‘N+1’ lane; ‘N’ indicates 20-mer amino primer and ‘N+1’ indicates the extended 21-mer product).

**Figure 1:**
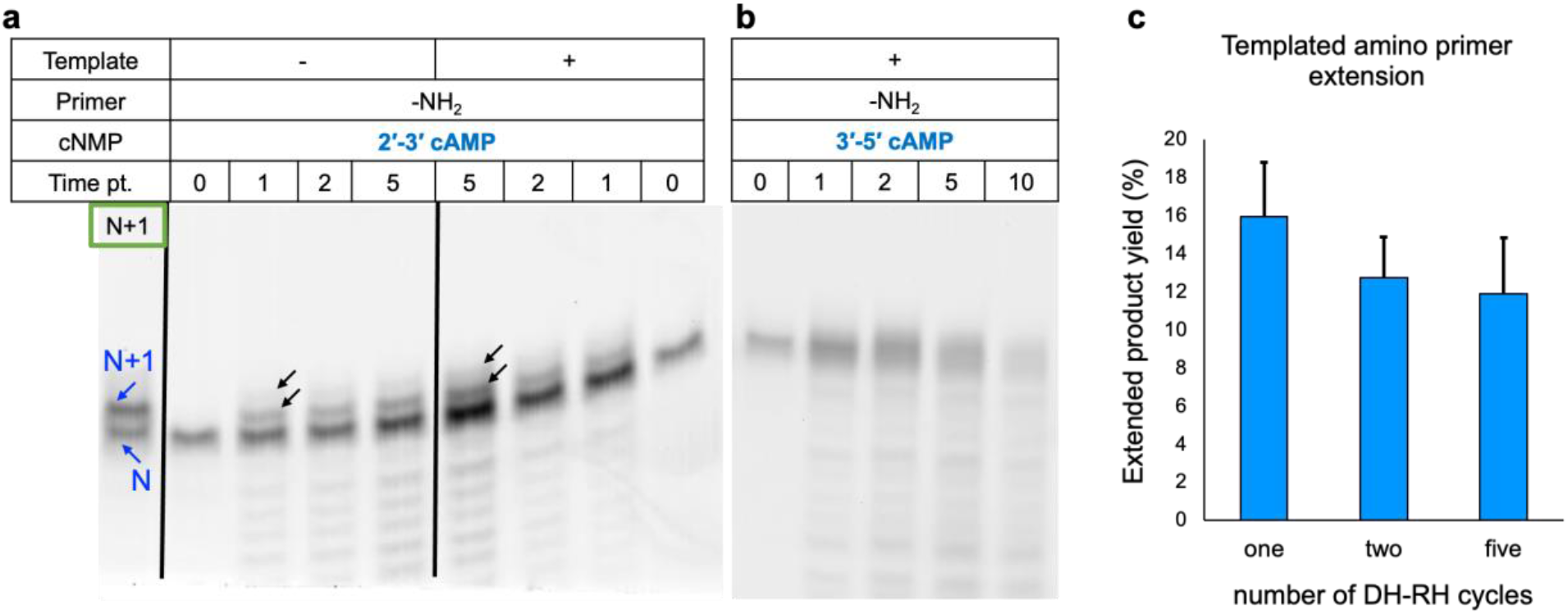
Extension of the NH_2_-primer using cAMP as the monomer over multiple cycles of DH-RH. The reactions were performed using 2′, 3′ cAMP (a) or 3′, 5′ cAMP (b), either in the absence (1a, left panel) or presence of template U (1a right panel and b), and analyzed after varying number of DH-RH cycles (cycle 0, cycle 1, cycle 2, cycle 5 and cycles 10). In the N+1 lane, blue arrows indicating ‘N’ and ‘N+1’ denote the control 20-mer RNA primer and the primer extended by one nucleotide (21-mer RNA), respectively. The black arrows indicate the extended products in the respective reactions showing up to two nucleotide additions as against the control lane. The black vertical lines have been used to demarcate two reaction sets that were run on the same gel. (c) Yield of the extended primer product in the reactions involving template-directed NH_2_-primer reaction, by analyzing denaturing gels using Image Quant v8.2. Y-axis shows the quantified yield percent of the total extended product (sum of both the extended bands), after different DH-RH cycles (viz., 1, 2 and 5) as indicated in the X-axis. The difference was found to be insignificant based on a two-tailed t-test. Error bars = s.d., N = 3.

At pH 2, the intensity of ‘N’ band decreased with increasing DH-RH cycles, with no extension observed (Fig. S3 and S8). This could be because of low stability of RNA under these harsh conditions (pH 2 and 90°C). However, at both pH 8 and 10, two new bands above the ‘N’ band, indicating the presence of species with higher molecular weight, were observed (Fig. 1a, S3 and S8). These new bands indicated the possibility of extension by one (N+1) and two nucleotides (N+2), respectively (Fig. 1a). In the case of pH 10, up to 9.4% of the extended product was observed after one DH-RH cycle. Nonetheless, the intensity of the bands corresponding to the extended products was observed to decrease significantly with increasing number of DH-RH cycles. Only 5.6 % of the extended product was observed after two cycles at pH 10, which further decreased to 3.8% after five DH-RH cycles. The reason behind this could be the instability of particularly the phosphodiester bond at alkaline pH. The template-directed reactions carried out at pH 8 gave optimal yield, with up to 15.9 % of primer being extended within one DH-RH cycle (Fig. 1c). The yield of the extended product was observed to decrease to 12.7 % and 11.9 % after two and five DH-RH cycles, respectively. Pertinently, this extended product was also observed in untemplated reactions (Fig. 1a). In the case of untemplated reactions, up to 12.4 % primer was observed to extend within one DH-RH cycle (Fig. S1a, S2a and 2b). Similar to the template-directed reactions, the yield of extended primer decreased to 10.6 % and 7.7 % after two and five DH-RH cycles, respectively. Given the aforementioned observations, all subsequent reactions were performed at pH 8.

As mentioned earlier, the two cyclic isomers possess different reactivity owing to their corresponding ring strains. Encouraged by the primer extension using 2′, 3′ cAMP, we next investigated the template-directed primer extension reaction using 3′, 5′ cAMP. All other reaction conditions (such as temperature, pH and DH-RH duration) were kept the same as that of the reactions that were done using 2′, 3′ cAMP. 1 mM 3′, 5′ cAMP was used as the rehydrating agent in these reactions. Given the intrinsic higher stability of 3′, 5′ cAMP (over 2′, 3′ cAMP), the reaction mixtures were subjected to ten DH-RH cycles to characterize observable reaction changes.

As observed in Figure 1b, with increasing number of DH-RH cycles, the intensity of the ‘N’ band decreased. However, no clear primer extension bands were observed even after ten DH-RH cycles (Fig. 1b). This could be potentially because of the higher ring stability of 3′, 5′ cAMP, which makes it comparatively less reactive. Moreover, our reaction conditions of high temperature (90 °C) and alkaline pH (8) results in relatively greater RNA primer degradation when experienced for longer periods (e.g., ten days), as is seen in Figure 1b. This further adds to the incompatibility of using 3′, 5′ cAMP, as it potentially requires longer duration to react and facilitate template-directed primer extension reactions.

### Primer extension using OH-primer and cyclic nucleotides under DH-RH conditions

After undertaking preliminary template-directed primer extension reactions using NH_2_-primer where the extended product is linked with a phosphoramidite bond, we also used OH-primer to check its propensity for extension using cNMPs. It is pertinent to mention that earlier attempts involving template-directed OH-primer extension using non-activated nucleotides, did not result in intact single nucleotide-based extensions (17). This could be because of the lesser nucleophilicity of the hydroxyl group when compared to an amino group. Nonetheless, we wanted to investigate the capability of cyclic nucleotides to extend OH-primer, which would result in the contemporary phosphodiester inter-nucleotide linkage. Towards this, reactions were performed using OH-primer and 2′, 3′ cAMP. All other reaction conditions (concentrations of reactants, pH, temperature and DH-RH duration) were kept the same as in the reactions that involved the use of NH_2_-primer.

Contrary to when non-activated nucleotides were used, two extension bands of the OH-primer were observed in both the untemplated as well as the template-directed reactions; similar to what was seen with the NH_2_-primer (Fig. 2a) reactions. Up to 12.9% of the extension product yield was observed after one DH-RH cycle in these reactions. Up to 12.6% and 11% of extension products persisted even after two and five DH-RH cycles, respectively (Fig. 2b). In the case of untemplated OH-primer extension reactions, the extended product yield was observed to be 8% within one DH-RH cycle (Fig. S1b, S2c and S2d). Upon comparison, up to 11% of the extended product was observed to persist in the presence of template even after five DH-RH cycles, as compared to only ∼5.3 % in the untemplated reactions. Upon comparison of the OH-primer reactions with NH_2_-primer reactions, the difference in the yield of the extended product was observed to be insignificant, irrespective of the presence or absence of the template. This was really encouraging as the extension with OH-primer results in a canonical phosphodiester linkage instead of phosphoramidite linkage (as is the case when using NH_2_-primer). Moreover, to our knowledge, this is the only reaction to date that has been shown to extend contemporary hydroxyl RNA primer, without the requirement of activating either the nucleotide or the primer.

**Figure 2:**
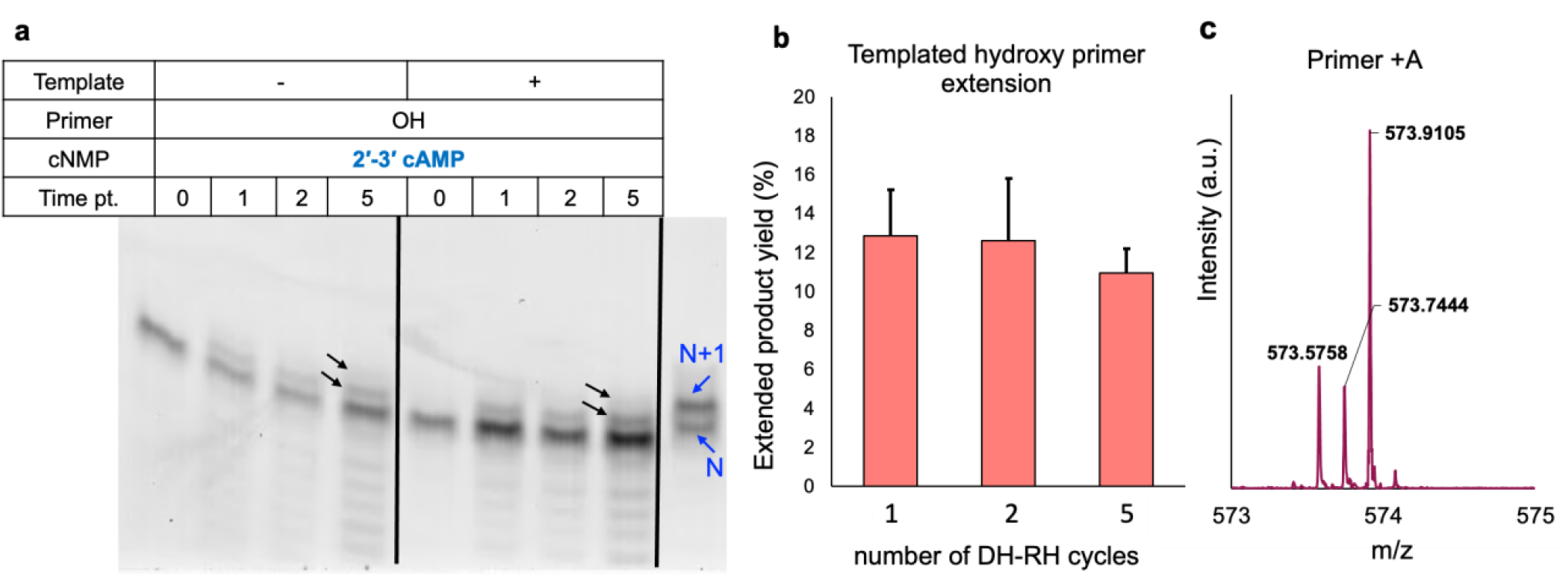
Extension of the OH-primer using 2′, 3′ cAMP as the monomer over repeated cycles of DH-RH. (a) The reactions were performed either in the absence (2a, left panel) or presence of template U (2a, right panel) and analyzed after varying number of DH-RH cycles (cycle 0, cycle 1, cycle 2 and cycle 5). In the N+1 lane, blue arrows indicating ‘N’ and ‘N+1’ denote the control 20-mer RNA primer and the extended 21-mer RNA, respectively. The black arrows indicate the extended product in the respective reaction mixtures. The black vertical lines have been used to demarcate two reaction sets that were run on the same gel. (b) Yield of the extended primer product with the OH-primer, obtained by analyzing the denaturing gels using Image Quant v8.2. Y-axis shows the quantified yield percent of the total extended product (sum of both the extended bands), after different DH-RH cycles (viz., one, two and five) as is indicated in the X-axis. The difference was found to be insignificant based on a two-tailed t-test. Error bars = s.d., N = 3. (c) Representative TOF-MS spectrum in ESI negative mode for cAMP reactions showing the extended 10-mer primer with the addition of AMP, resulting in a 11-mer primer [M] with m/z corresponding to [M-6H]^6-^ = 573.9105.

This is exciting as the use of cNMPs would have readily facilitated uphill oligomerization and templated replication reactions, without the need for invoking any activation chemistries in the prebiotic scenario. Additionally, similar to the reactions with NH_2_-primer, no extension bands were observed when 3′, 5′ cAMP was used instead of 2′, 3′ cAMP (Fig. S5).

In order to confirm whether the extended products seen in the aforementioned contexts are indeed intact nucleotide additions, we performed the same reaction but with a 10-mer OH-primer (detailed in Methods section), to facilitate a comprehensive analysis of the reaction mixture using LC-MS (detailed in the Methods section). The presence of m/z corresponding to intact nucleotide additions to yield ‘N+1’ (Fig. 2c) and ‘N+2’ (Table S1) within ± 5 ppm error, confirmed beyond doubt that the primer extensions in reactions using 2′, 3′ cAMP indeed are extended products with intact informational moieties. This is contrary to scenarios that led to the sugar-phosphate backbone extension but resulted in abasic sites that were a consequence of deglycosylation of the informational entities, resulting in molecules that would lack the capacity for hydrogen bonding-based functionalities (17, 23).

### Effect of lipids on the cNMP-based primer extension reactions under DH-RH conditions

Encouraged by the addition of two intact nucleotides in the aforementioned primer extension reactions involving 2′, 3′ cNMP, we investigated the effect of lipids on these reactions. Previously, the presence of lipids has been demonstrated to protect the molecules involved in the nonenzymatic oligomerization and template-directed primer extension reactions, against hydrolysis. Further, their presence also imparts an identity to the encapsulated system that form instantaneously post rehydration in such scenarios (17, 23, 25). Given this, primer extension reactions were also performed in the presence of 5 mM POPC for the template-directed as well as untemplated reactions, using both OH-primer and NH_2_-primer.

As shown in Figure 4 (panels a-d), extension till two bases (‘N+1’ and ‘N+2’) was observed in all the reactions. The presence of lipids was observed to enhance the yield of extended products in both the templated as well as untemplated reactions (Fig. 4e, 4f and S4). In the case of POPC-assisted template-directed OH-primer reactions, yield of the extended product enhanced from 12.9% (in the absence of POPC) to 18.4 % within one DH-RH cycle, and 15.5% of the product persisted even after five DH-RH cycles (Fig. 4e and 3b). Similarly, in the untemplated reactions, the presence of POPC led to an increase in the product yield to 13.3% (as compared to 8% in the absence of POC) within one DH-RH cycle, wherein 9.6% of it remained even after five DH-RH cycles. In the NH_2_-primer containing POPC-assisted reactions, the yield enhanced to 13.6% (12.4% in the absence of POPC) and 21.6% (15.9% in the absence of POPC) in untemplated and template-directed reactions, respectively (Fig. 4f and 3a). Up to 9.5% (in untemplated) and 19.8% (in template-directed) of the product persisted even after five DH-RH cycles in the presence of POPC, whereas only 7.7 % and 11.9 % was observed in its absence (in untemplated and template-directed reactions, respectively). This increase in the extended primer yield in the presence of POPC could be because of its protecting effect towards the substrates (primer as well as cyclic nucleotides) against hydrolysis. This, in turn, would have enhanced the availability of the reacting substrates, thereby leading to an enhancement in the product yield.

**Figure 4:**
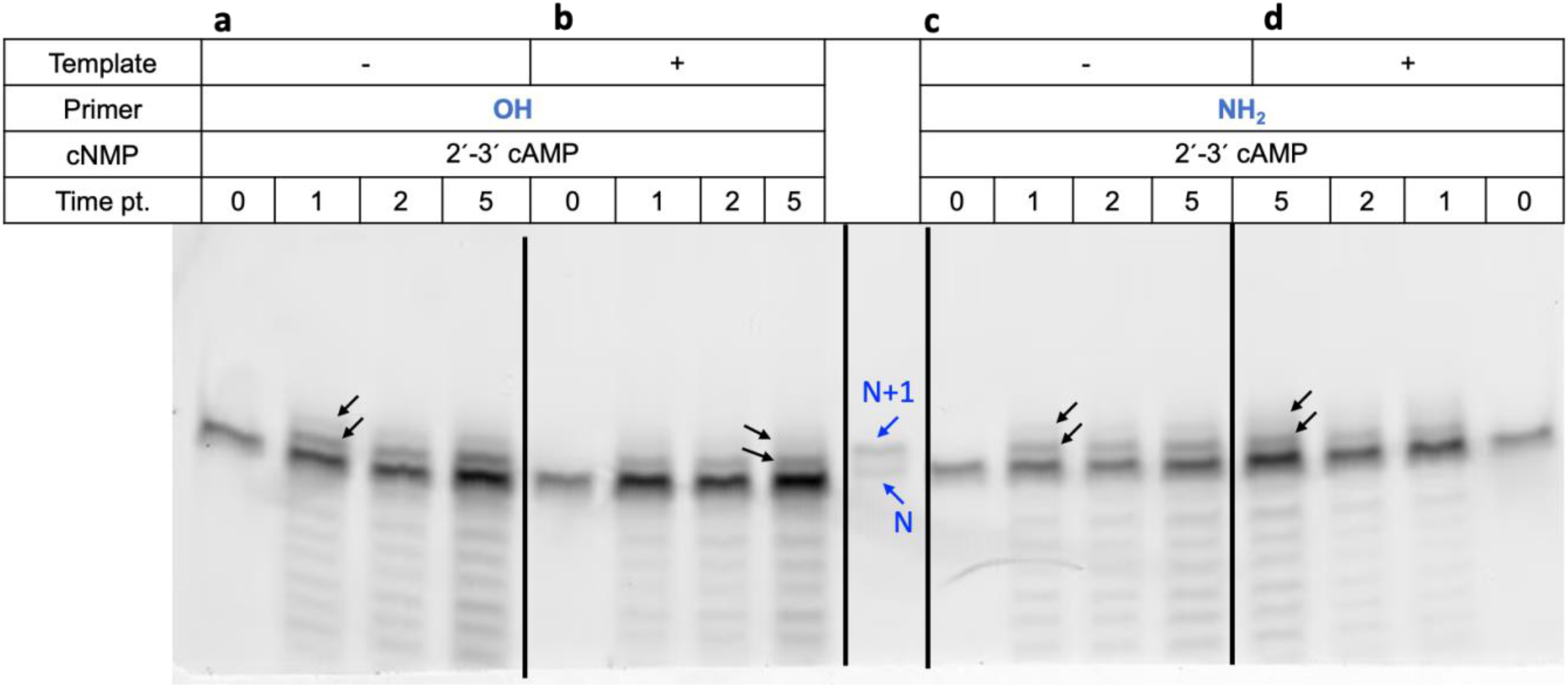
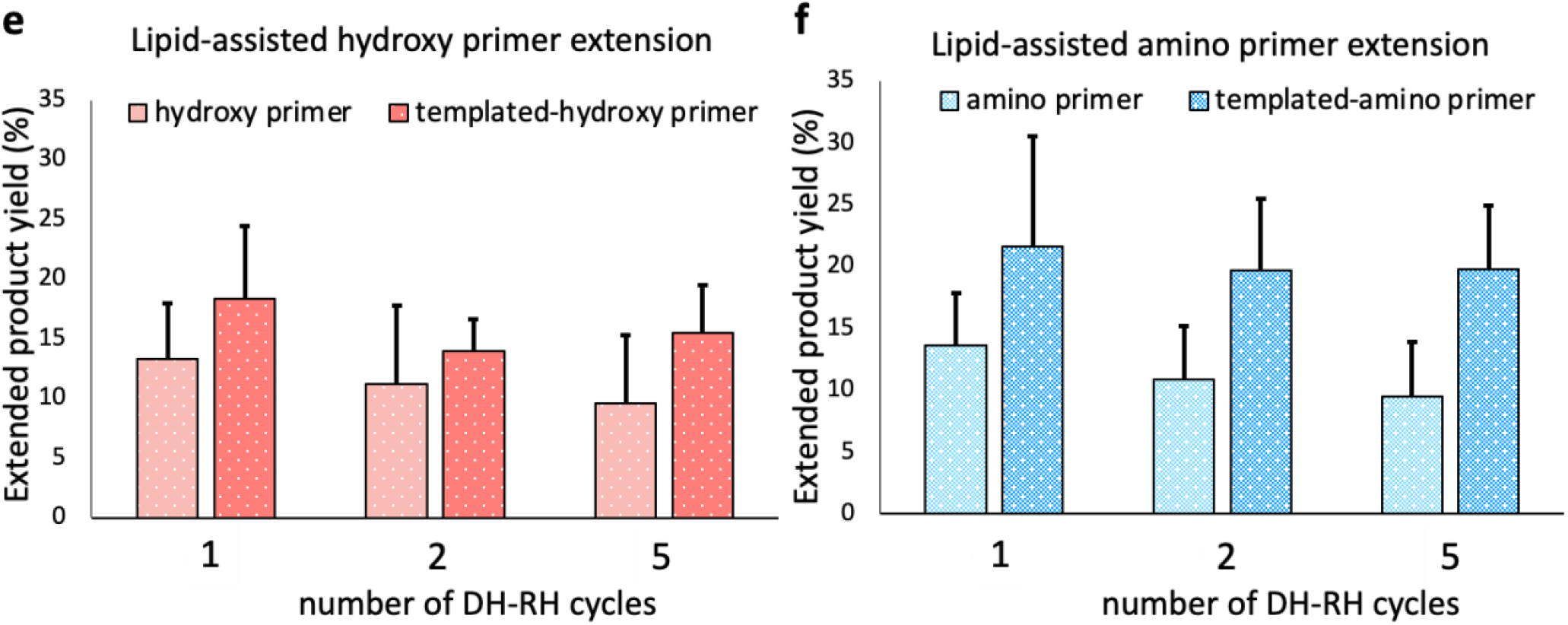
Effect of POPC on RNA primer extension. Reactions were performed using template U, 2′, 3′ cAMP, and either NH_2_-primer or OH-primer, over repeated cycles of DH-RH (i.e., cycle 0, cycle 1, cycle 2 and cycle 5). The black arrows indicate the extended product. In the N+1 lane, ‘N’ indicates the control 20-mer RNA primer while ‘N + 1’ indicates extension of the primer by one nucleotide. The black vertical lines in the above two panels have been used to demarcate two reaction sets that were run on the same gel. Panels e and f show the quantified total yield (%) of the extended primer in the lipid-assisted reactions of OH-primer (panel e) and NH_2_-primer (panel f), respectively. Yields were quantified for both the untemplated and template-directed reactions (as depicted in the legend), after varying the number of DH-RH cycles (x-axis). The difference was found to be insignificant based on a two-tailed t-test. Error bars = s.d., N=3.

Importantly, as mentioned earlier, we investigated the effect of prebiotically realistic hybrid compartments on our enzyme-free template-directed primer extension reactions. Towards this, prebiotically relevant composite compartments comprising POPC along with equimolar concentrations of either oleic acid (OA) or glycerol monooleate (GMO), were investigated (see Methods for details). Similar to the POPC-assisted reactions, both POPC:OA and POPC:GMO systems were observed to enhance the yield of the extended product when compared to the reactions without any lipid in them (Fig. 5, Table 1). Nonetheless, the difference between the yield of extended products in the presence of any of the membrane systems was found to be insignificant (based on a two-tailed t-test; Table S5 and S6) These results signify that all the lipid compositions evaluated seemed to impart protection to the extended products against hydrolysis as the amount of extended product did not decrease significantly (based on a two-tailed t-test) even after five DH-RH cycles

**Figure 5:**
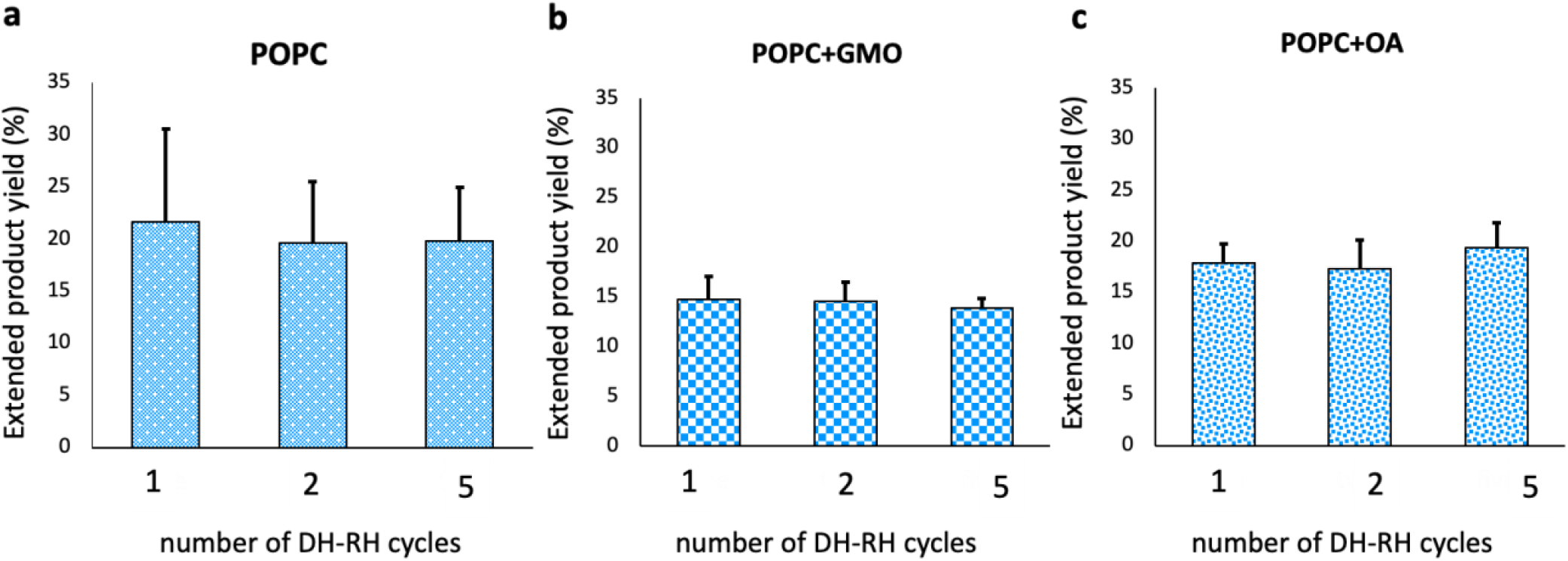
Effect of different lipids on lipid-assisted RNA primer extension. Reactions were performed using template U, 2′, 3′ cAMP and NH_2_-primer, in the presence of 5 mM lipid, over repeated cycles of DH-RH, as indicated by X-axis. Different panels show the quantified yield (%) of extended product in the presence of compartments with different lipid compositions, viz. pure POPC (a), 1:1 POPC:GMO (b) and 1:1 POPC:OA (c). The difference was found to be insignificant based on a two-tailed t-test. Error bars = s.d., N=3.

**Table 1:** The yield (%) of the extended product in the presence of different lipid systems i.e., pure POPC and two hybrid membrane systems (1:1 POPC:GMO and 1:1 POPC:OA) over repeated DH-RH cycles.

| **Lipid-assisted template-directed primer extension using cAMP** |  |  |  |
| --- | --- | --- | --- |
|  | **number of DH-RH cycles** |  |  |
|  | **1** | **2** | **5** |
| **POPC** | 21.61 ± 8.91 | 19.67 ± 5.83 | 19.80 ± 5.14 |
| **POPC + GMO** | 14.73 ± 2.32 | 14.58 ± 1.92 | 13.92 ± 0.92 |
| **POPC + OA** | 17.83 ± 1.93 | 17.31 ± 2.85 | 19.38 ± 2.48 |

**Table 2:** The yield (%) of the extended product in the presence and absence of POPC using 2′, 3′ cCMP and with NH_2_-primer or OH-primer over repeated DH-RH cycles.

| Template-directed primer extension using cCMP |  |  |  |
| --- | --- | --- | --- |
|  | number of DH-RH cycles |  |  |
|  | 1 | 2 | 5 |
| **OH-primer** | 3.47 ± 0.5 | 2.49 ± 0.25 | 1.32 ± 0.2 |
| **NH<sub>2</sub>-primer** | 3.36 ± 0.04 | 2.85 ± 0.22 | 1.95 ± 0.09 |
| **POPC-assisted OH-primer** | 7.04 ± 0.74 | 5.19 ± 1.11 | 6.37 ± 1.03 |
| **POPC-assisted NH<sub>2</sub>-primer** | 11.55 ± 2.36 | 9.63 ± 1.52 | 8.12 ± 1.35 |

### Primer extension using a pyrimidine-based cyclic nucleotides (cCMP) under DH-RH conditions

Our results showed template-directed primer extension using cAMP as the incoming nucleotide, against uracil as the templating base. Importantly, purines are known to stack better and potentially have a better tendency to undergo concentration-dependent condensation reactions. Similar results were also observed in the nonenzymatic oligomerizations reactions, where the yield of oligomers in cCMP reactions were lower than that of cAMP (25). Therefore, we examined how the presence of a pyrimidine-based cNMP i.e., cCMP, would influence these reactions as compared to purine-based cAMP. Towards this, we performed template-directed primer extensions using cyclic 2′, 3′ cCMP (Fig.6) against template G. The reactions were performed with both OH-primer and NH_2_-primer.

**Figure 6:**
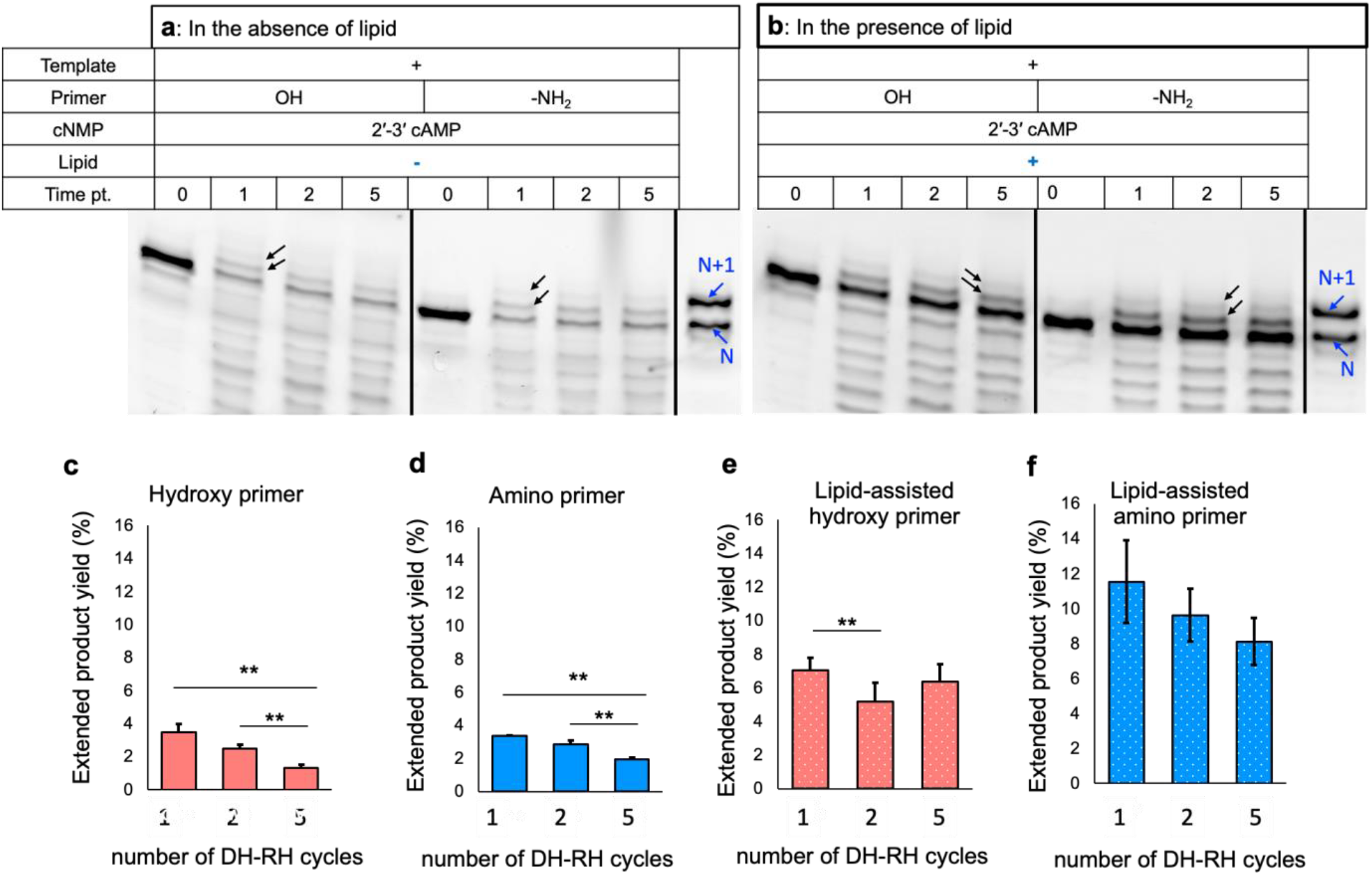
Primer extension using cCMP under multiple DH-RH cycles and effect of lipids on these reactions. Reactions were performed using template G, 2′, 3′ cCMP and with NH_2_-primer or OH-primer, over repeated DH-RH cycles in the absence (a) or in the presence of 5 mM POPC (b). The black arrows indicate the extended products. In the N+1 lane, ‘N’ indicates the control 20-mer RNA primer while ‘N + 1’ indicates extension of the primer by one nucleotide. The black vertical lines in the above two panels have been used to demarcate two reaction sets that were run on the same gel. N=3. Panels c to f show the quantified total yield (%) of the extended primer in the absence of POPC (c and d) and POPC-assisted reactions (e and f), for OH-primer and NH_2_-primer, respectively. Yields were quantified for both untemplated and templated reactions (as depicted in the legend) after varying number of DH-RH cycles (x-axis). The ** indicates a significant change with a p-value<0.01 based on a two-tailed t-test. The difference was found to be insignificant for the remaining bars. Error bars = s.d., N=3.

As observed in Figure 6a, the primer extension with up to two nucleotide additions was observed for both of these primers when analyzed using PAGE (Fig. 6a). The intact nucleotide additions in the primer extension reactions (involving a 10-mer primer) were further confirmed using LC-MS. The m/z corresponding to ‘N+1’ and ‘N+2’ was observed within less than 5 ppm error (Table S2). Upon quantification, 3.5% and 3.4% of the extended product was observed within one DH-RH cycle in the case of OH-primer and NH_2_-primer, respectively (Fig. 6a, 6c and 6d). With an increase in the number of DH-RH cycles to five, the yield of the extended product decreased significantly (based on a two-tailed t-test, Table S7) for both the OH-primer (1.32%) and NH_2_-primer (1.9%). The reason behind this could be the instability of RNA under our reaction conditions i.e., 90°C and pH 8 for five DH-RH cycles.

As alluded to earlier, the presence of lipids under such scenarios has been shown to impart protection against hydrolysis. In order to investigate the effect of lipids on template-directed replication, the extension reactions were performed using cCMP against template G in the presence of POPC. As observed in Figure 6b, two extension bands were seen in the lipid-assisted extension of both OH-primer and NH_2_-primer within one DH-RH cycle. Both the extension bands persisted till five DH-RH cycles in these reactions. Upon quantification, the yield of extended product in these POPC-assisted reactions was observed to be significantly higher (based on a two-tailed t-test, Table S10) than in the reactions without lipid. In the case of POPC-assisted OH-primer reactions, 7.0 % of the extended product was observed within one DH-RH cycle (Fig. 6e) as compared to just 3.5 % in the corresponding reactions without lipid. Moreover, the decrease in the extended product yield after five DH-RH cycles was found to be insignificant (based on a two-tailed t-test, Table S9) in POPC-assisted OH-primer reactions. Conversely, in the reactions without lipids, with increasing number of DH-RH cycles, the yield of extended product significantly decreased (Table S7).

In the case of POPC-assisted NH_2_-primer reactions, 11.6 % of extended primer was observed within one DH-RH cycle for NH_2_-primer (Fig. 6f). This was significantly higher when compared to the 3.4 % yield obtained in the absence of POPC (Fig. 6d). Moreover, the decrease in the extended product yield after five DH-RH cycles was found to be insignificant (based on a two-tailed t-test) in the POPC-assisted NH_2_-primer reactions (Fig. 6f). These results emphasize the fact that the yield and stability (against hydrolysis) of the extended primer was enhanced significantly based on a two-tailed t-test (Table S10) in the lipid-assisted reactions.

## Discussion

Enzyme-free oligomerization of RNA and its propagation, are central to the RNA World Hypothesis (2). Previous studies in this context have predominantly employed activated nucleotides to study these phenomena (4, 38–44). However, the availability of activated nucleotides in significant amounts on the prebiotic Earth is debatable (25, 45). In this context, a related study demonstrated template-directed primer extension reactions using non-activated 5′-NMPs under acidic terrestrial geothermal conditions i.e., acidic pH, high temperature and DH-RH cycles (17). However, systematic characterization of these extended products indicated abasic sites due to the cleavage of the glycosidic bond especially because of the acidic pH (23, 24). Since cNMPs are intrinsically reactive due to the ring strain present in them, they do not require harsh conditions such as acidic pH for undergoing oligomerization reactions (46). In an earlier study, we demonstrated the nonenzymatic oligomerization of these cNMPs under high temperature, alkaline conditions (pH 8), using repetitive dry-wet cycles (25). In this study, we demonstrate the enzyme-free template-directed primer extension using cNMPs, under the same conditions. Our results show primer extension happening, with up to two nucleotide additions in all the reactions investigated, using 2′, 3′cNMP under alkaline conditions (pH 8 and 10). Nonetheless, alkaline conditions are known to be challenging for RNA stability as it leads to phosphodiester bond cleavage. This explains the optimal yield of the extended product at pH 8. However, similar reactions with 3′, 5′ cNMP did not yield in any extension products under our experimental conditions. This could be due to their relatively higher stability towards hydrolysis when compared to 2′, 3′ cAMP (25, 46). Also, 3′, 5′ cNMP contains a six-membered ring when compared to a five-membered ring in 2′, 3′ cNMP. This makes it comparatively less reactive due to the lower ring strain that it experiences (8.9 kcal/mol) when compared to the five-membered ring of 2′,3′ cAMP (12 kcal/mol, 45).

Pertinently, the extension of primer was observed with both OH and NH_2_-primers, for both cAMP and cCMP containing reactions. This signifies the generalizability of primer extension using cNMPs without the imminent need for any external activation for facilitating these reactions. Upon comparing the yield of the extended product, the cAMP containing template-directed reactions were observed to be significantly higher yielding as compared to the corresponding cCMP reactions (Table S11). In the case of template-directed reactions using OH-primer, the difference in cAMP and cCMP reactions, after one and two DH-RH cycles, was found to be insignificant based on a two-tailed t-test (Table S11). After five DH-RH cycles, the extended product in cAMP was found to be significantly (Table S11) higher. In template-directed NH_2_-primer extension reactions, after one and two DH-RH cycles, the extended product was significantly higher in cAMP (Table S11). However, by five DH-RH cycles, the difference was found to be insignificant (Table S11). In lipid-assisted reactions, the primer extension was observed to occur in the presence of POPC as well as prebiotically relevant hybrid membrane systems (POPC+GMO and POPC+OA), highlighting the compatibility of these reactions with model protocellular membranes. In the case of cCMP-based reactions, the presence of PL was observed to significantly increase the yield of the extended product. This could be because of the capability of lipid to impart protection to RNA as well as cNMPs against hydrolysis (25).

Most previous studies have employed activated nucleotides to study primer extension reactions, and these are facilitated at ambient temperatures (6, 38, 43, 47–49). Therefore, one of the crucial challenges faced by enzyme-free template-directed replication reactions under these ambient conditions, is “strand separation” (39). This is essential for the replicated duplex strands to separate, in order to allow for multiple rounds of information propagation. Towards this, few studies that involved the use of activated nucleotides, have suggested workarounds including the use of viscous solvent, pH change or incorporating backbone heterogeneity in order to overcome this problem (39, 50, 51). In our study, the template-directed primer extension reactions occur under prebiotically relevant conditions of high temperature (90°C), and under geochemically pertinent DH-RH cycles. These recurrent DH-RH fluctuations at 90°C can, in principle, facilitate the annealing of primer and template to form duplex during the “dehydrated” phase and assist in strand separation in the “rehydrated” phase due to high temperature aqueous conditions. Additionally, 2′, 3′ cNMP involve intramolecular phosphodiester bonds, which upon oligomerization and primer extension, result in extended products with inherently mixed intermolecular phosphodiester linkages. Such random backbone heterogeneity in the duplexes is known to result in their melting at lower temperatures (50), further facilitating in effective strand separation. Given these direct implications for RNA propagation on the early Earth, our study underlines the importance of facilitating template-directed primer extension reactions, using intrinsically active cNMPs under prebiotically pertinent terrestrial geothermal conditions. Further, our results also highlight the intrinsic capability of environmental fluctuations such as DH-RH cycles, in catalyzing nonenzymatic information propagation on the early Earth, thus renewing our understanding about how oligomerization and propagation of RNA would have occurred under realistic prebiotic conditions, during the emergence of a putative RNA World.

## Author contributions

S.D. and S.R. designed the experiments. S.S. performed and analyzed the reactions with varying lipid compositions. S.D. performed the remainder of the experiments. S.D. and S.R. analyzed the data. S.D., S.S. and S.R. wrote the manuscript.

## Acknowledgements

This research was supported by grants from the Science and Engineering Research Board (SERB), Department of Science and Technology, Govt. of India [EMR/2015/000434], the Department of Biotechnology, Govt. of India [BT/PR19201/BRB/10/1532/2016] and IISER Pune. S.D. and S.S. acknowledge CSIR, Govt. of India and IISER Pune, respectively, for their fellowship.

