## Supplementary Information for "Nonenzymatic template-directed replication using 2′-3′ cyclic nucleotides under wet-dry cycles"

### Supporting Information

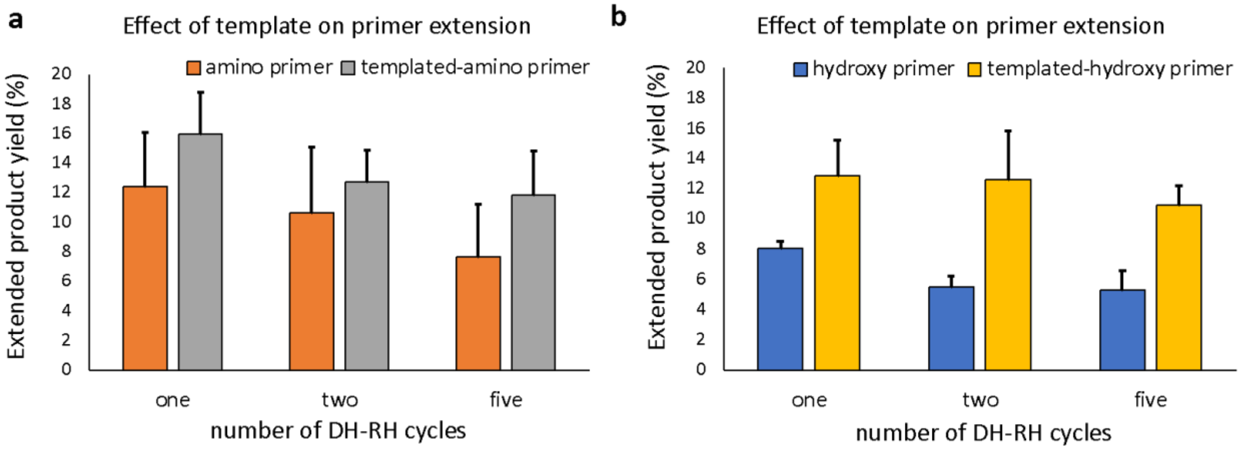


**Figure S1: Effect of presence of template U on the extension of RNA primer using 2′, 3′ cAMP.** The reactions were performed using either $NH_2$-primer (a) or OH-primer (b), over repeated cycles of DH-RH (i.e., one, two and five). Y-axis shows the total yield (%) of the extended product and x-axis shows the number of DH-RH cycles. Error bars = s.d., N = 3.

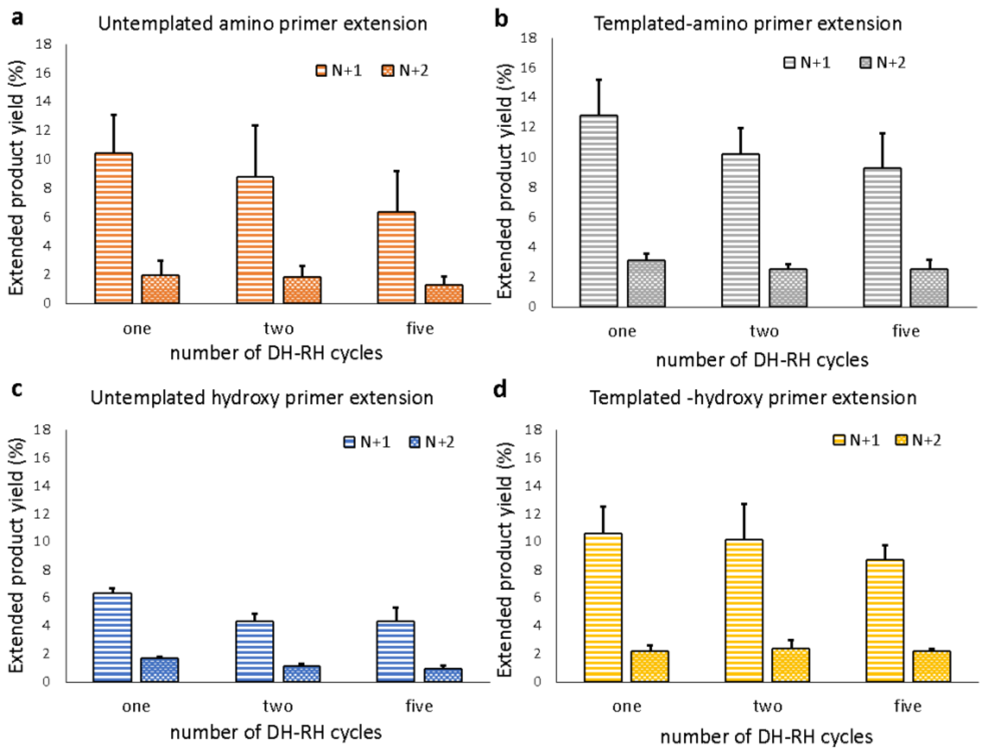


**Figure S2: Effect of presence of template U on the yield of ‘N+1’ and ‘N+2’ extended products using 2′, 3′ cAMP.** a) and b): The reactions were performed using $NH_2$-primer in the absence (a) or presence of template-U (b), over repeated cycles of DH-RH (i.e., one, two and five). c) and d): The reactions were performed using OH-primer in the absence (c) or presence of template-U (d), over repeated cycles of DH-RH (i.e., one, two and five). Y-axis shows the yield (%) of the extended product (‘N+1’ and ‘N+2’) and x-axis shows the number of DH-RH cycles. Error bars = s.d., N = 3.

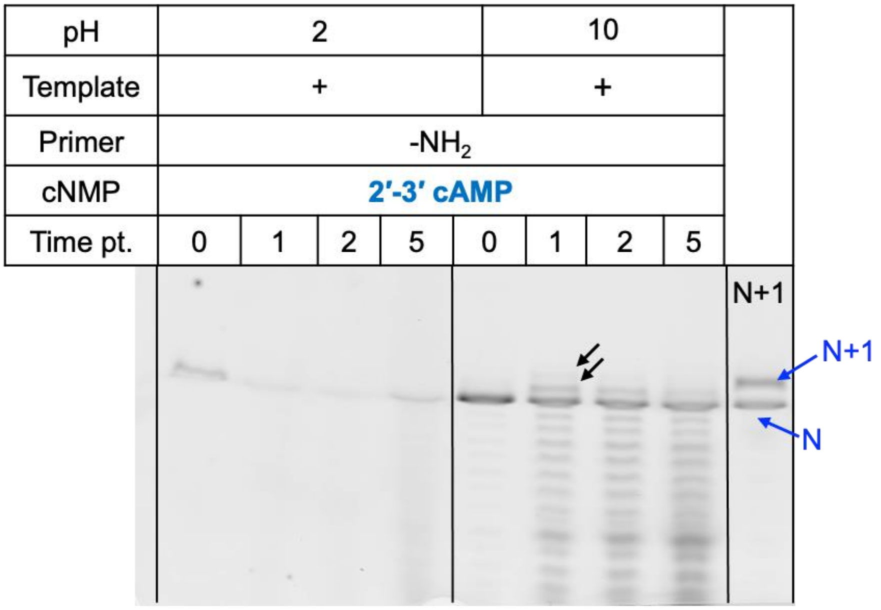


**Figure S3: Effect of varying pH on the template-directed primer extension reaction. The** reactions were performed using template-U, $NH_2$-primer and 2′, 3′ cAMP as the monomer, over multiple DH-RH cycles (cycle 0, cycle 1, cycle 2 and cycle 5) at pH 2 (left panel) and 10 (right panel). In the N+1 lane, blue arrows indicating 'N' and 'N+1' denote the control 20-mer RNA primer and the primer extended by one nucleotide (21-mer RNA), respectively. The black arrows indicate the extended products in the reactions at pH 10, showing up to two nucleotide additions as against the control lane. The black vertical lines have been used to demarcate two reaction sets that were run on the same gel. All the reactions were repeated at least thrice (N = 3).

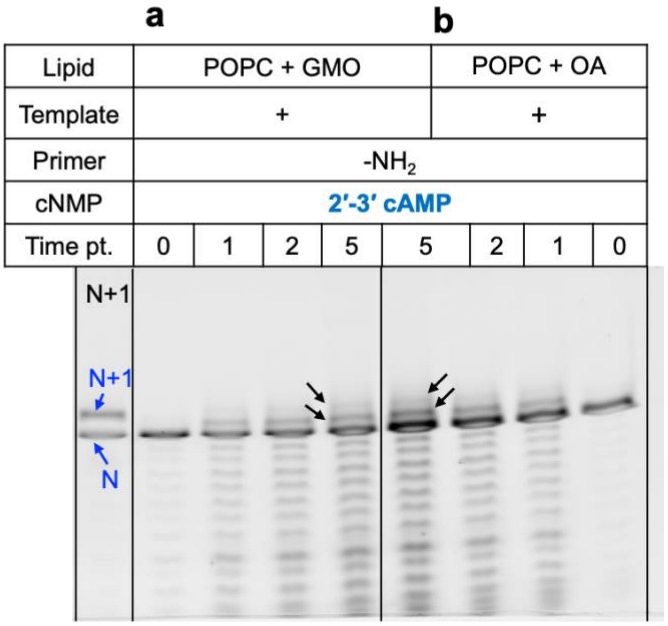


**Figure S4: Effect of varying lipid composition on the lipid-assisted primer extension reaction.** Reactions were performed using template-U, $NH_2$-primer and 2′, 3′ cAMP as the monomer, over multiple DH-RH cycles (cycle 0, cycle 1, cycle 2 and cycle 5) in the presence of 5 mM lipid. The different lipid compartments used are as follows, (a) 1:1 POPC:GMO and (b)1:1 POPC:OA. In the N+1 lane, blue arrows indicating 'N' and 'N+1' denote the control 20-mer RNA primer and the primer extended by one nucleotide (21-mer RNA), respectively. The black arrows indicate the extended products in the reactions, showing up to two nucleotide additions as against the control lane. The black vertical lines have been used to demarcate two reaction sets that were run on the same gel. All the reactions were repeated at least thrice (N = 3).

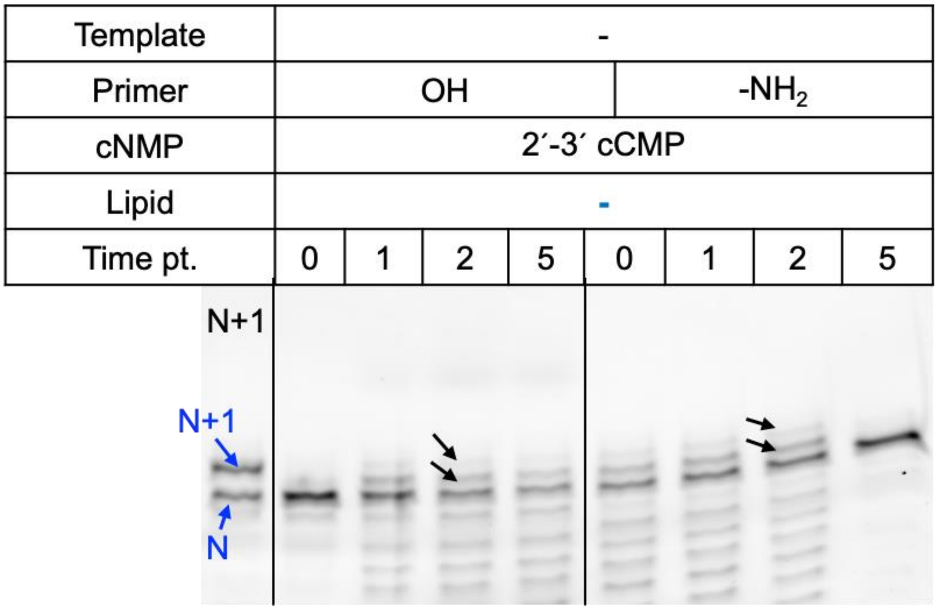


**Figure S5: The untemplated primer extension reactions using 2′, 3′ cCMP.** Reactions were performed using either OH-primer (left panel) or $NH_2$-primer (right panel), over multiple DH-RH cycles (cycle 0, cycle 1, cycle 2 and cycle 5). In the N+1 lane, blue arrows indicating ‘N’ and ‘N+1’ denote the control 20-mer RNA primer and the primer extended by one nucleotide (21-mer RNA), respectively. The black arrows indicate the extended products in the reactions, showing up to two nucleotide additions as against the control lane. The black vertical lines have been used to demarcate two reaction sets that were run on the same gel. All the reactions were repeated at least thrice (N = 3).

**The uncropped original images of all the gels in the main article and supporting information.** The images were acquired using Amersham Typhoon Biomolecular imager (GE Healthcare) at 550 PMT and 100-micron resolution setting, using the Cy3 (532 nm) excitation laser. The gel images were subsequently processed using ImageQuant v8.2 software for quantification of the relevant bands.

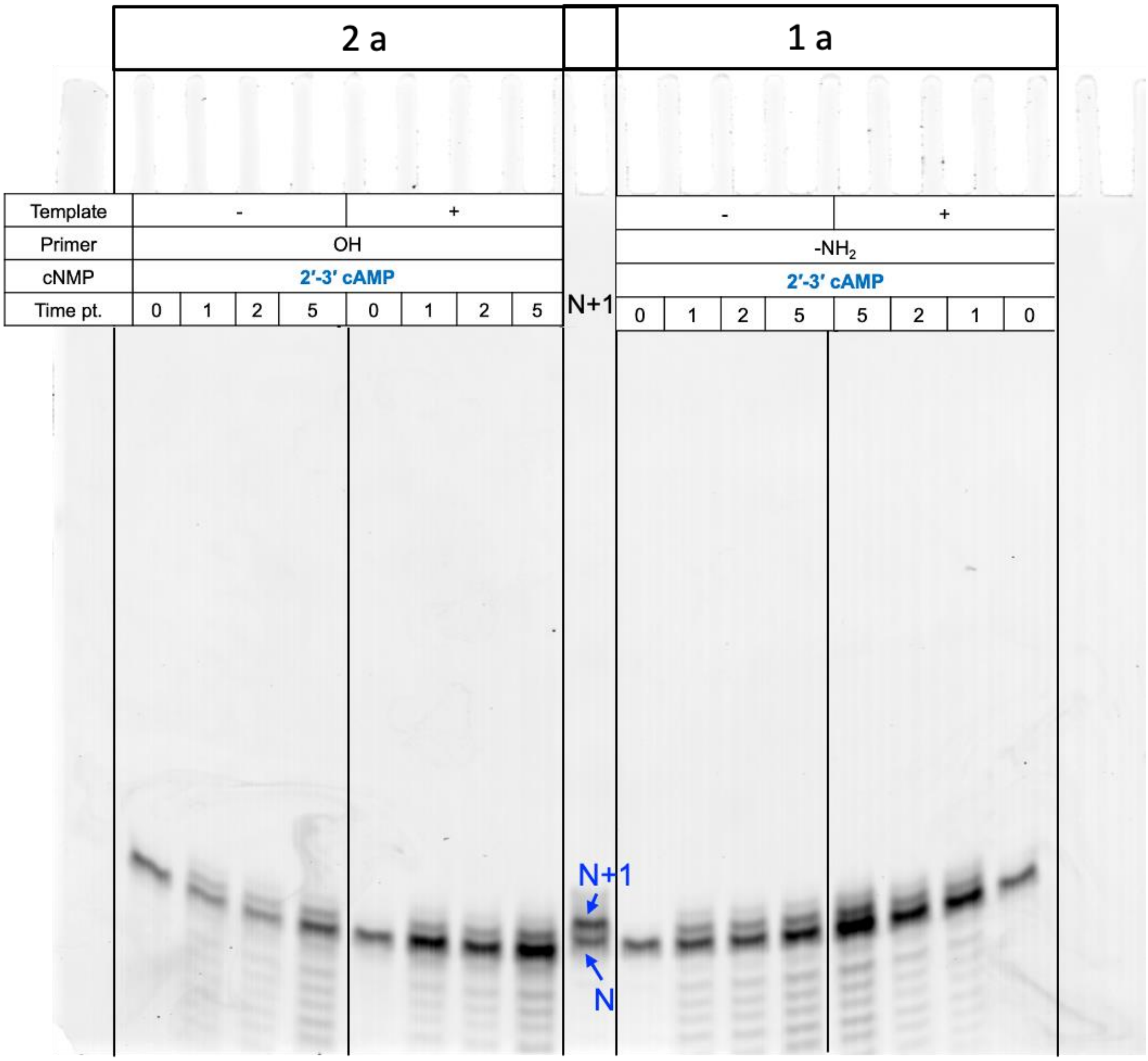


**Figure S6: The uncropped original gel image for the main paper Figures 2a and 1a, respectively.** 2a) The reactions were performed using 2′, 3′ cAMP and OH-primer either in the absence (2a, left panel) or presence of template U (2a, right panel) and analyzed after varying number of DH-RH cycles (cycle 0, cycle 1, cycle 2 and cycle 5). 1a) The reactions were performed using 2′, 3′ cAMP and $NH_2$-primer either in the absence (1a, left panel) or presence of template U (1a, right panel) and analyzed after varying number of DH-RH cycles (cycle 0, cycle 1, cycle 2 and cycle 5). In the N+1 lane, blue arrows indicating 'N' and 'N+1' denote the control 20-mer RNA primer and the primer extended by one nucleotide (21-mer RNA), respectively. Y-axis shows the yield (%) of the extended product ('N+1' and 'N+2') and x-axis shows the number of DH-RH cycles. All the reactions were repeated at least thrice (N = 3).

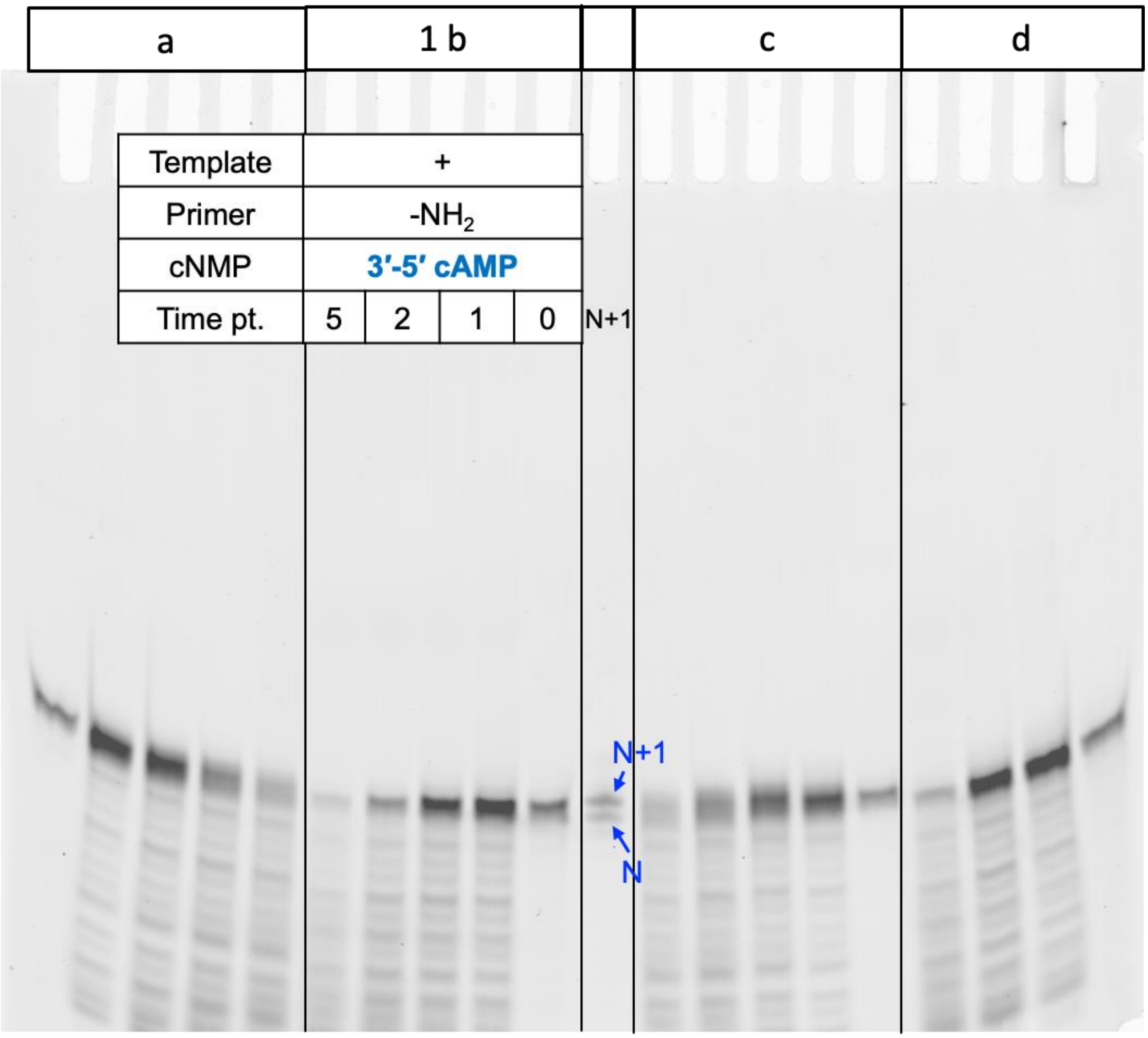


**Figure S7: The uncropped original gel image for the main paper Figure 1b.** Extension of the $NH_2$-primer using 3′, 5′ cAMP as the monomer in the presence of template-U, over multiple cycles of DH-RH (cycle 0, cycle 1, cycle 2 and cycle 5). Panel a shows the extension of the OH-primer using 3′, 5′ cAMP as the monomer in the presence of template-U, over multiple cycles of DH-RH cycles. Panels c and d show the untemplated control reactions of the OH-primer and $NH_2$-primer, respectively, over multiple DH-RH cycles using 3′, 5′ cAMP. No extension bands were observed in any of the reactions 3′, 5′ cAMP containing reactions. In the N+1 lane, blue arrows indicating 'N' and 'N+1' denote the control 20-mer RNA primer and the primer extended by one nucleotide (21-mer RNA), respectively. All the reactions were repeated at least thrice (N = 3).

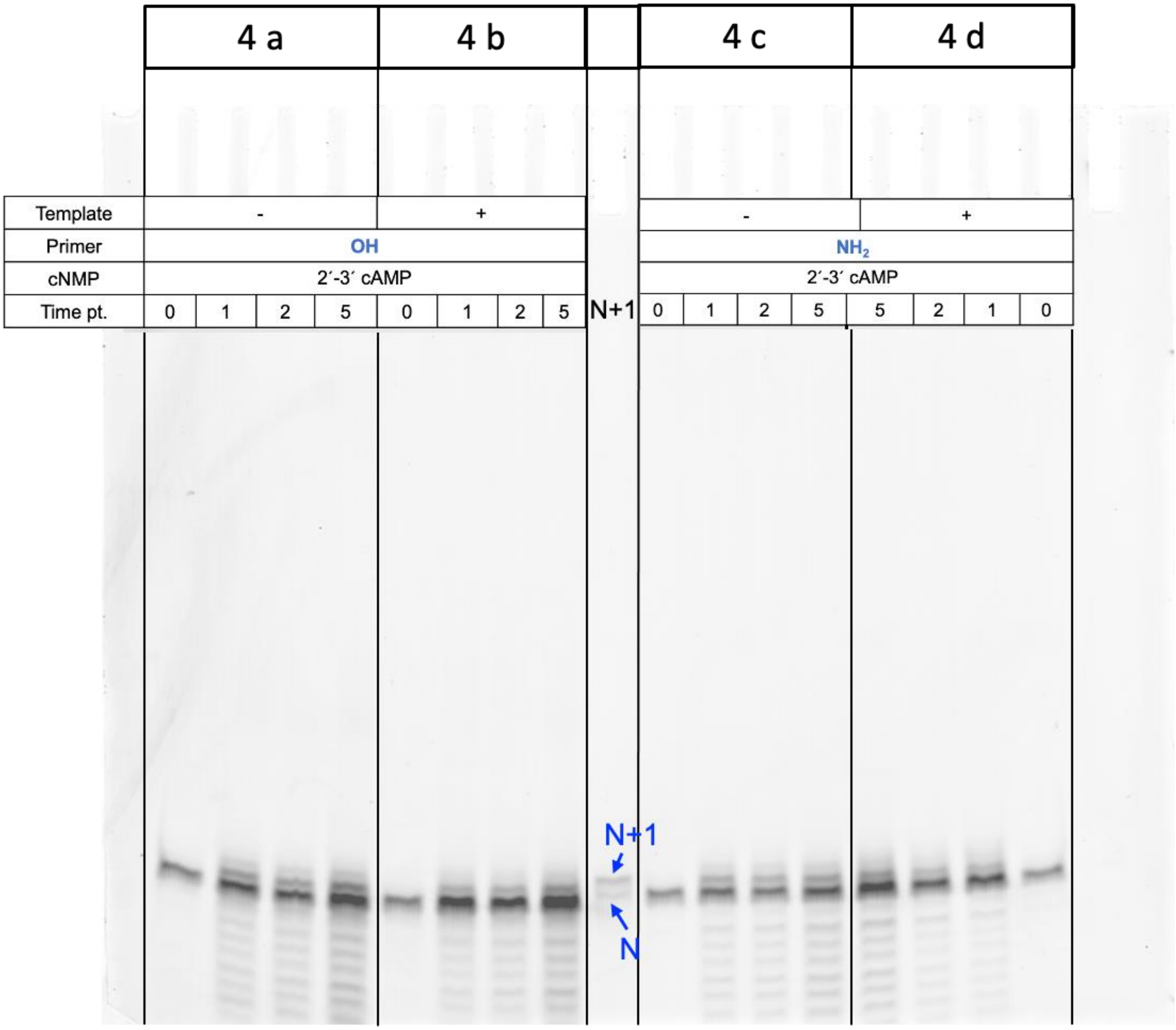


**Figure S8: The uncropped original gel image for the main paper Figure 4.** Effect of POPC on RNA primer extension. Reactions were performed in the presence and absence of template U using 2′, 3′ cAMP, using either $NH_2$-primer or OH-primer, over repeated cycles of DH-RH (i.e., cycle 0, cycle 1, cycle 2 and cycle 5). In the N+1 lane, blue arrows indicating ‘N’ and ‘N+1’ denote the control 20-mer RNA primer and the primer extended by one nucleotide (21-mer RNA), respectively. All the reactions were replicated at least thrice (N = 3).

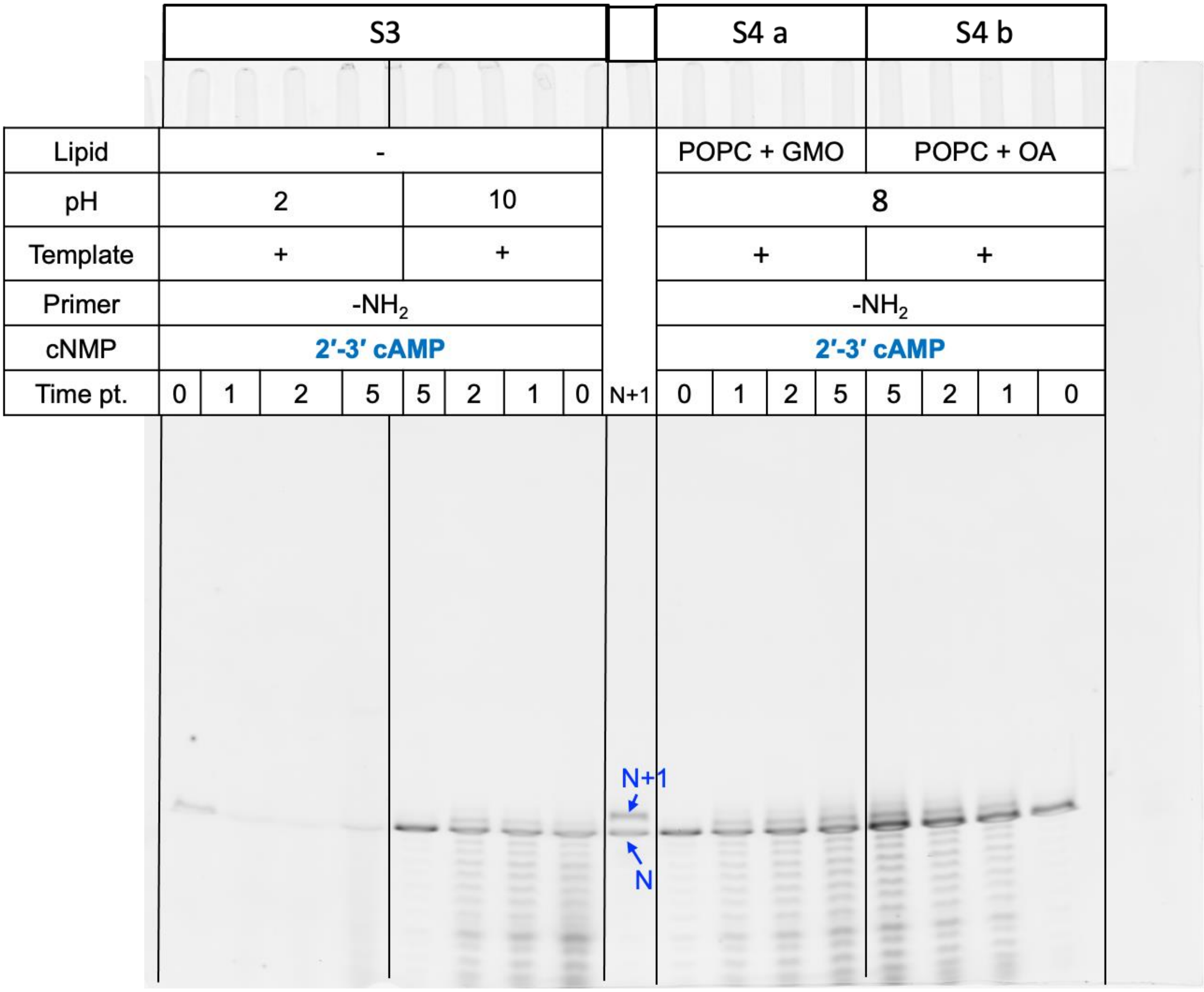


**Figure S9: The uncropped original gel image for supplementary Figures S3 and S4.** S3) Effect of varying pH (2 and 10) on the extension reactions. Reactions were performed using template-U, $NH_2$-primer and 2′, 3′ cAMP as the monomer, over multiple DH-RH cycles (cycle 0, cycle 1, cycle 2 and cycle 5) at pH 2 (left panel) and 10 (right panel), respectively. S4) Effect of varying lipid composition on the lipid-assisted primer extension reactions. Reactions were performed using template-U, $NH_2$-primer and 2′, 3′ cAMP as the monomer, over multiple DH-RH cycles (cycle 0, cycle 1, cycle 2 and cycle 5) in the presence of 5 mM lipid. Different lipid compartments used were (a) 1:1 POPC:GMO and (b) 1:1 POPC:OA. In the N+1 lane, blue arrows indicating ‘N’ and ‘N+1’ denote the control 20-mer RNA primer and the primer extended by one nucleotide (21-mer RNA), respectively. All the reactions were replicated at least thrice (N = 3).

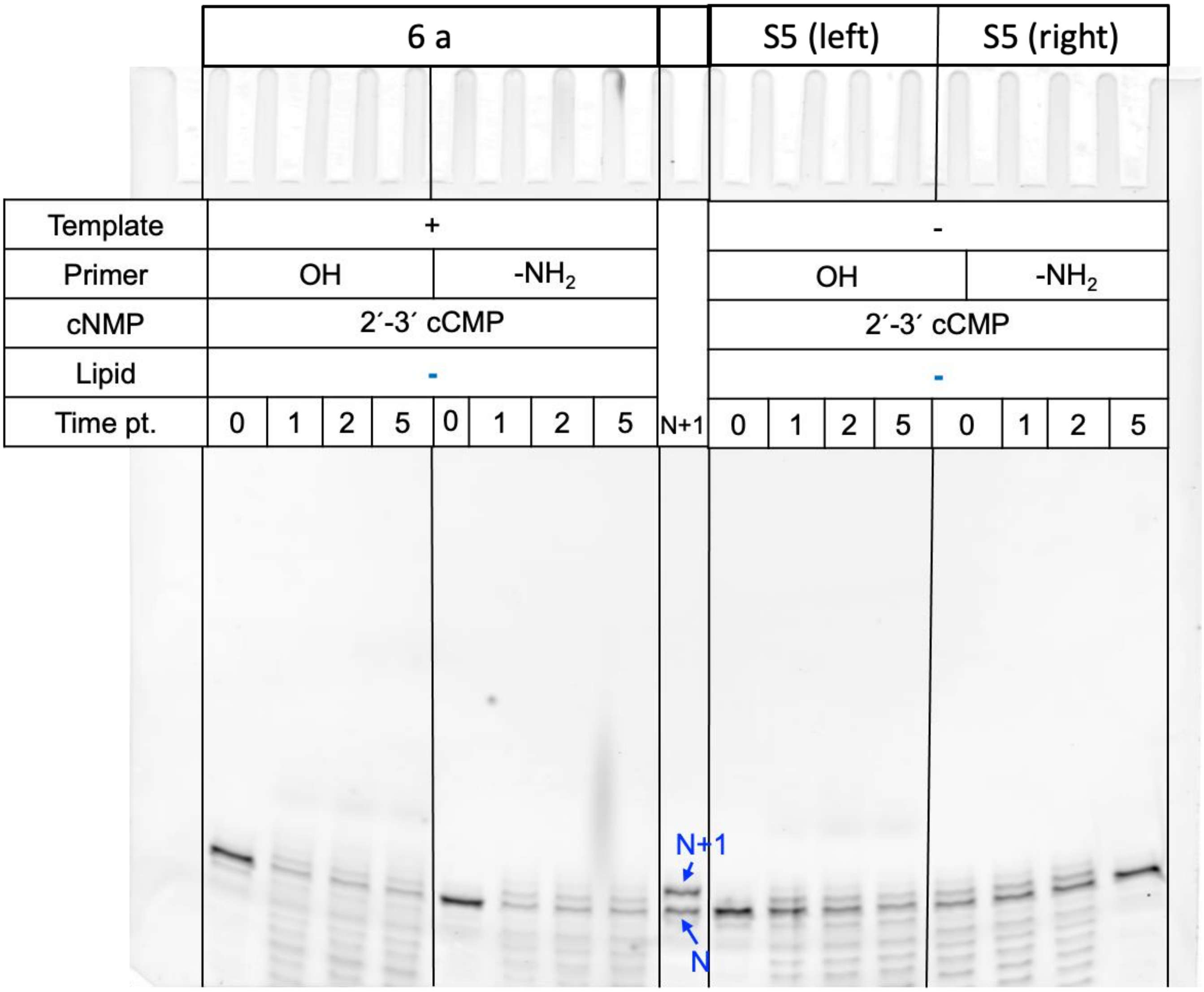


**Figure S10: The uncropped gel image for the main paper Figures 6a and S5.** Reactions were performed using template G, 2′, 3′ cCMP and with either $NH_2$-primer or OH-primer, over repeated DH-RH cycles (i.e., cycle 0, cycle 1, cycle 2 and cycle 5). Panels b and c show the untemplated control reactions of OH-primer and $NH_2$-primer, respectively, over multiple DH-RH cycles using 2′, 3′ cCMP. In the N+1 lane, blue arrows indicating ‘N’ and ‘N+1’ denote the control 20-mer RNA primer and the primer extended by one nucleotide (21-mer RNA), respectively. All the reactions were replicated at least thrice (N = 3).

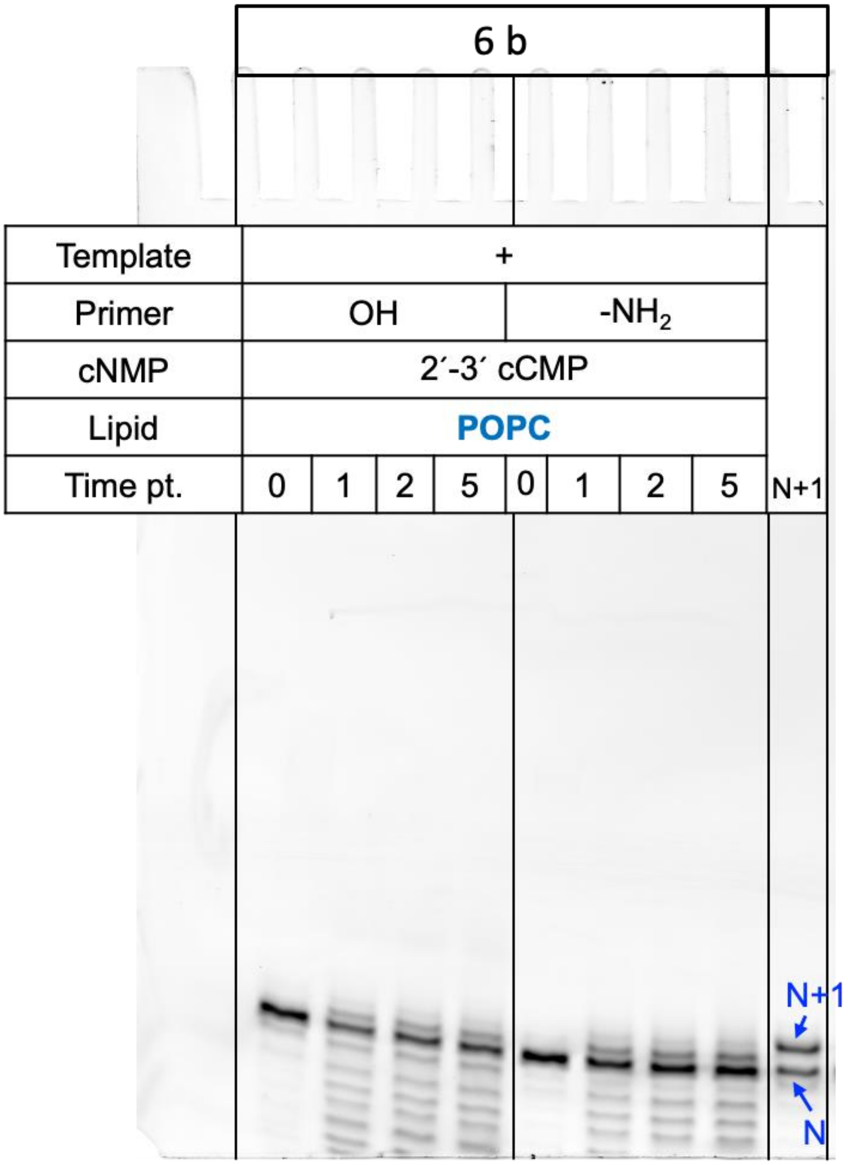


**Figure S11: The uncropped gel image for the main paper Figure 6b.** Reactions were performed using template G, 2′, 3′ cCMP and with either $NH_2$-primer or OH-primer in the presence of 5 mM POPC, over repeated DH-RH cycles (i.e., cycle 0, cycle 1, cycle 2 and cycle 5). In the N+1 lane, blue arrows indicating 'N' and 'N+1' denote the control 20-mer RNA primer and the primer extended by one nucleotide (21-mer RNA), respectively. All the reactions were repeated at least thrice (N = 3).

**Table S1: Masses observed during LC-MS characterization of primer extension reactions using 2′, 3′ cAMP, along with calculated parts per million (ppm) error. Masses for one nucleotide and two nucleotide extensions on to the 10-mer OH-primer have been indicated as ‘N+1’ and ‘N+2’, respectively.**

| 2′-3′ cAMP | | | | |
|---|---|---|---|---|
| | **z** | **m/z (calculated)** | **m/z (observed)** | **ppm error** |
| **N + 1** | 3 | 1148.1635 | 1148.1562 | -6.3 |
| | 4 | 861.6233 | 861.6223 | -1.16 |
| | 5 | 688.6959 | 688.6979 | 2.97 |
| | 6 | **573.5781** | **573.5782** | 0.31 |
| **N +2** | 8 | 471.0633 | 471.0636 | 0.57 |
| | 6 | 628.4202 | 628.4203 | 0.18 |
| | 5 | 754.3057 | 754.3070 | 1.79 |
| | 4 | 943.3848 | 943.3850 | 0.20 |
| | 3 | 1258.1821 | 1258.1803 | -1.42 |

**Table S2: Masses observed during LC-MS characterization of primer extension reactions using 2′, 3′ cCMP, along with calculated parts per million (ppm) error. Masses for one nucleotide and two nucleotide extensions on to the 10-mer OH-primer have been indicated as ‘N+1’ and ‘N+2’, respectively.**

| 2′-3′ cCMP | | | | |
|---|---|---|---|---|
| | **z** | **m/z (calculated)** | **m/z (observed)** | **ppm error** |
| **N +1** | 2 | 1710.7433 | 1710.73666 | -3.89 |
| | 3 | 1140.1598 | 1140.15784 | -1.7 |
| | 4 | 854.8680 | 854.8696 | 1.87 |
| | 7 | 488.064321 | 488.064456 | 0.28 |
| **N +2** | 1 | 3728.5385 | 3728.5309 | -2.03 |
| | 2 | 1863.7656 | 1863.7699 | 2.32 |
| | 3 | 1242.1746 | 1242.1729 | -1.39 |
| | 5 | 744.9019 | 744.9059 | 5.34 |
| | 7 | 531.7850 | 531.7852 | 0.36 |
| | 4 | 931.3792 | 931.3779 | -1.36 |
| | 6 | 620.5837 | 620.5846 | 1.39 |

### Statistical Analysis

**Table S3: Comparison of the primer extension in untemplated and template-directed reactions, using 2′, 3′ cAMP with either $NH_2$-primer or OH-primer, between different DH-RH cycles (cycle 1, 2 and 5) using two-tailed type 2 t-test. (Significant p-value is highlighted with a green background). N = 3.**

| **2′, 3′ cAMP** | | | | | | | | |
|---|---|---|---|---|---|---|---|---|
| Cycles being compared | Untemplated | | | | Template-directed | | | |
| | $NH_2$-primer | | OH-primer | | $NH_2$-primer | | OH-primer | |
| | p-value | | p-value | | p-value | | | p-value |
| 1 with 2 | 0.77 | ns | 0.042 | * | 0.41 | ns | 0.96 | ns |
| 2 with 5 | 0.63 | ns | 0.90 | ns | 0.82 | ns | 0.68 | ns |
| 1 with 5 | 0.41 | ns | 0.12 | ns | 0.36 | ns | 0.55 | ns |

**Table S4: Comparison of the primer extension in untemplated vs template-directed reactions, using 2′, 3′ cAMP with either $NH_2$-primer or OH-primer across various DH-RH cycles between the two reactions, using two-tailed type 2 t-test. (The difference was found to be insignificant). N = 3.**

| **2′, 3′ cAMP** | | | | | | | | |
|---|---|---|---|---|---|---|---|---|
| Cycles being compared | Untemplated $NH_2$-primer vs OH-primer | | Untemplated vs template-directed $NH_2$-primer | | Untemplated vs template-directed OH-primer | | Template-directed $NH_2$-primer vs OH-primer | |
| | p-value | | p-value | | p-value | | | p-value |
| 1 with 1 | 0.30 | ns | 0.48 | ns | 0.08 | ns | 0.54 | ns |
| 2 with 2 | 0.31 | ns | 0.66 | ns | 0.07 | ns | 0.98 | ns |
| 5 with 5 | 0.56 | ns | 0.40 | ns | 0.06 | ns | 0.85 | ns |

**Table S5: Comparison of the primer extension reactions in without lipid reactions vs lipid-assisted reactions, using 2′, 3′ cAMP with either $NH_2$-primer or OH-primer, in the presence or absence of template U, across various DH-RH cycles, using two-tailed type 2 t-test. (The difference was found to be insignificant). N = 3.**

| **2′, 3′ cAMP** | | | | | | | | |
|---|---|---|---|---|---|---|---|---|
| Cycles being compared | Without lipid vs lipid-assisted | | | | | | | |
| | Untemplated | | | | Template-directed | | | |
| | $NH_2$-primer | | OH-primer | | $NH_2$-primer | | OH-primer | |
| | p-value | | p-value | | p-value | | | p-value |
| 1 with 1 | 0.83 | ns | 0.13 | ns | 0.38 | ns | 0.38 | ns |
| 2 with 2 | 0.97 | ns | 0.21 | ns | 0.16 | ns | 0.75 | ns |
| 5 with 5 | 0.74 | ns | 0.30 | ns | 0.18 | ns | 0.27 | ns |

**Table S6: Comparison of different lipids on lipid-assisted RNA primer extension reactions using 2′, 3′ cAMP and $NH_2$-primer, across various DH-RH cycles between the two reactions, using two-tailed type 2 t-test. (Significant p-value is highlighted with a green background). N = 3.**

| **Template-directed $NH_2$-primer extension using 2′, 3′ cAMP** | | | | | | |
|---|---|---|---|---|---|---|
| Cycles being compared | POPC : GMO vs POPC : OA | | POPC : GMO vs POPC | | POPC : OA vs POPC | |
| | p-value | | p-value | | p-value | |
| 1 with 1 | 0.15 | ns | 0.26 | ns | 0.498 | ns |
| 2 with 2 | 0.24 | ns | 0.23 | ns | 0.57 | ns |
| 5 with 5 | 0.02 | * | 0.126 | ns | 0.90 | ns |

**Table S7: Comparison of template-directed primer extension, using 2′, 3′ cCMP with either $NH_2$-primer or OH-primer, between different DH-RH cycles, using two-tailed type 2 t-test. (Significant p-value is highlighted with a green background). N = 3.**

| 2′, 3′ cCMP | | | | |
|---|---|---|---|---|
| Cycles being compared | Template-directed | | | |
| | OH-primer | | $NH_2$-primer | |
| | p-value | | p-value | |
| 1 with 2 | 0.13 | ns | 0.08 | ns |
| 2 with 5 | 0.04 | * | 0.03 | * |
| 1 with 5 | 0.03 | * | 0.002 | ** |

**Table S8: Comparison of template-directed primer extension, using $NH_2$-primer vs OH-primer in 2′, 3′ cCMP containing reactions, across various DH-RH cycles between the two reactions, using two-tailed type 2 t-test. (The difference was found to be insignificant). N = 3.**

| 2′, 3′ cCMP | | |
|---|---|---|
| Cycles being compared | Template-directed | |
| | $NH_2$-primer vs OH-primer | |
| | p-value | |
| 1 with 1 | 0.80 | ns |
| 2 with 2 | 0.27 | ns |
| 5 with 5 | 0.06 | ns |

**Table S9: Comparison of lipid-assisted template-directed primer extension, using $NH_2$-primer vs OH-primer in 2′, 3′ cCMP containing reactions, across various DH-RH cycles between the two reactions, using two-tailed type 2 t-test. (Significant p-value is highlighted with a green background). N = 3.**

| **2′, 3′ cCMP** | | |
|---|---|---|
| Cycles being compared | Lipid-assisted | |
| | Template-directed | |
| | $NH_2$-primer vs OH-primer | |
| | p-value | |
| 1 with 1 | 0.01 | * |
| 2 with 2 | 0.003 | ** |
| 5 with 5 | 0.08 | ns |

**Table S10: Comparison of template-directed, without lipid vs lipid-assisted primer extension, using either $NH_2$-primer or OH-primer in 2′, 3′ cCMP containing reactions, across various DH-RH cycles between the two reactions, using two-tailed type 2 t-test. (Significant p-value is highlighted with a green background). N = 3.**

| **2′, 3′ cCMP** | | | | |
|---|---|---|---|---|
| Cycles being compared | Template-directed | | | |
| | Without lipid vs lipid-assisted | | | |
| | OH-primer | | $NH_2$-primer | |
| 1 with 1 | 0.004 | ** | 0.009 | ** |
| 2 with 2 | 0.03 | * | 0.004 | ** |
| 5 with 5 | 0.003 | ** | 0.004 | ** |

**Table S11: Comparison of template-directed RNA primer extension using 2′, 3′ cAMP vs 2′, 3′ cCMP in either without lipid or lipid-assisted reactions, across various DH-RH cycles between the two reactions, using two-tailed type 2 t-test. (Significant p-value is highlighted with a green background). N = 3.**

| **2′, 3′ cAMP vs 2′, 3′ cCMP** | | | | | | | | |
|---|---|---|---|---|---|---|---|---|
| Template -directed | | | | | | | | |
| Cycles being compared | Without lipid | | | | Lipid-assited | | | |
| | $NH_2$-primer | | OH-primer | | $NH_2$-primer | | OH-primer | |
| | p-value | | p-value | | p-value | | p-value | |
| 1 with 1 | 0.04 | * | 0.06 | ns | 0.077 | ns | 0.13 | * |
| 2 with 2 | 0.037 | * | 0.09 | ns | 0.02 | * | 0.003 | ** |
| 5 with 5 | 0.09 | ns | 0.016 | * | 0.009 | ** | 0.008 | ** |
